# Principal and Independent Genomic Components of Brain Structure and Function

**DOI:** 10.1101/2022.07.13.499912

**Authors:** Lennart M. Oblong, Sourena Soheili-Nezhad, Nicolò Trevisan, Yingjie Shi, Christian F. Beckmann, Emma Sprooten

## Abstract

**Introduction:** The highly polygenic and pleiotropic nature of behavioural traits, psychiatric disorders, and structural and functional brain phenotypes complicate mechanistic interpretation of related genome-wide association study (GWAS) signals, such that the underlying causal biological processes remain obscure. We propose the novel method of genomic principal and independent component analysis (PCA, ICA) to decompose a large set of univariate GWAS statistics of multimodal brain traits into more interpretable latent genomic components. Here we introduce this new method and evaluate its various analytic parameters and reproducibility across independent samples.

**Methods:** Two releases of GWAS summary statistics from the UK biobank (UKB), with 11,086 and 22,138 participants respectively, were retrieved from the Oxford BIG-40 server. GWAS summary statistics were clumped resulting in n=165,364 single nucleotide polymorphisms (SNP) and m=2,240 imaging derived phenotypes (IDPs). Both genome-wide beta-values and their corresponding, standard-error scaled z-values were decomposed using multivariate exploratory linear optimised decomposition into independent components (MELODIC). We evaluated variance explained at multiple dimensions up to 200. We tested the reproducibility of output of dimensions 5, 10, 25, and 50 by computing Pearson’s correlation between component loadings, and Fisher Exact tests on overlap of the top SNP loadings across samples. Reproducibility statistics of the original raw and z-transformed univariate GWAS served as benchmarks. We also inspected the clustering of genomic components across neuroimaging modalities using t-SNE.

**Results:** The first five PCs derived from z-transformed GWAS captured 31.9% of the variance across SNP effect sizes, while 200 PCs increased the variance explained to 79.6%. Reproducibility of 10-dimensional PCs and ICs showed the best balance between model complexity and robustness, and variance explained (PCs: |r_z-max_|=0.33, |r_raw-max_|=0.30; ICs: |r_z-max_|=0.23, |r_raw-max_|=0.19), with decreasing model stability and reproducibility at higher dimensions. Both genomic PC and IC reproducibilities improved substantially relative to mean univariate GWAS reproducibility up to a dimension of 10. Genomic components clustered along neuroimaging modalities.

**Conclusion:** Our results indicate that these novel methods of genomic ICA and PCA decompose genetic effects on IDPs from raw GWAS statistics with high reproducibility by taking advantage of the inherent pleiotropic patterns. These findings encourage further applications of genomic ICA and PCA as fully data-driven methods to effectively reduce the dimensionality, enhance the signal to noise ratio, and improve interpretability of high-dimensional multi-trait genome-wide analyses.

## Introduction

Individual differences in brain structure and function are determined by complex biological mechanisms that remain largely unknown. Measures of human brain structure and function, as assessed through magnetic resonance imaging (MRI), are heritable (Elliott et al., 2018; Eyler et al., 2011; Glahn et al., 2010; Grasby et al., 2020; Jahanshad et al., 2013; McKay et al., 2014; Pizzagalli et al., 2020; Smith et al., 2021; Sprooten et al., 2014; Winkler et al., 2010). This indicates that genetic research on human brain structure and function may help to better understand the causal biological pathways that subserve the brain. Genome-wide association studies (GWAS) have emerged as a popular and powerful tool for estimating the effects of common genetic variants on behavioural traits, psychiatric disorders, and structural and functional brain phenotypes. This development is facilitated by the abundance of big genotypic and phenotypic data made available in biobanks and multi-center consortia, and showed that many features of the brain are polygenic, meaning that they are influenced by many common genetic variants in concert (Elliott et al., 2018; Hibar et al., 2017; Smith et al., 2021). Similar to multifactorial behavioural and psychiatric phenotypes (Wendt et al., 2020), the highly polygenic nature of these traits coupled with pleiotropic effects complicate interpretation of GWAS in terms of clear underlying causal biological processes (Watanabe et al., 2019; Matoba et al., 2022). Exploiting the high dimensional and pleiotropic nature of brain MRI-GWAS, we test the reproducibility of a novel, multivariate method that can help to translate the GWAS signal of multiple traits into more interpretable factors.

Generally, GWAS output consists of a large matrix which contains the mass-univariate statistics of millions of single nucleotide polymorphisms (SNPs) in relation to the trait investigated, thereby quantifying the genome-wide associations of common SNPs with a given trait. For most brain related traits this signal is highly polygenic, indicating the combined involvement of thousands of genes, each of which individually has a small effect (Elliott et al., 2018; Smith et al., 2021). The SNPs may impact on one or more levels of the causal biological processes subserving a trait, including DNA methylation, gene expression, protein synthesis, cellular functioning, ‘house-keeping’ mechanisms within the neuronal microenvironment, system-level brain morphology and functioning, and interactions with the environment. This long route of biological and molecular activity from genome to measured phenotype leads to uncertainty regarding the causal mechanisms underlying the GWAS signal. Thus, GWAS output reflects the final endpoints of a ‘mixture’ of many intermediate biological processes and molecular pathways. Groups of SNPs may share an involvement in the same biological pathways, which can be reflected in their covariation of effect sizes across different brain traits. Therefore, extracting hidden factors that capture shared SNP effects across brain traits may be insightful. These factors could provide novel insight into the shared biological mechanisms across brain structures, tissues and/or imaging modalities. We propose genomic independent component analysis (ICA) and genomic principal component analysis (PCA) applied to the GWAS summary statistics of thousands of structural and functional brain imaging derived phenotypes (IDPs), to identify hidden (i.e. latent) genomic factors influencing brain structure and function.

PCA and ICA are powerful decomposition methods to reduce the dimensionality of a large number of observations into fewer, often more interpretable components that capture covariation patterns across observations (Smith et al., 2020). PCA finds orthogonal components in the data that capture variance consecutively, with the first principal component (PC) capturing maximum variance, and the second and subsequent ones capturing variances orthogonal to the previous ones. Thus, in the context of present GWAS data, PCA will capture the maximum amount of shared genetic variance per IDP in as few components as possible. ICA on the other hand is an unsupervised source separation method that decomposes a complex signal into its constituent maximally independent parts, assuming a linear combination of (non-gaussian distributed) signal, structured noise, and gaussian distributed stochastic noise. ICA thus maximises independence between the components while allowing the component weights to be non-orthogonal if needed, which makes it more suitable to recover distinct sources of signal from noisy data that can be mixed within and across the initial variables (in our case SNPs) (Comon, 1994). Thus, compared to PCA, ICA captures variance less efficiently, but is designed to ‘unmix’ a complex signal into its constituents or sources, which become more visible and interpretable as a consequence (Hyvärinen & Oja, 2000). If we interpret current high-dimensional MRI-GWAS data as a signal composed of multiple underlying generative mechanisms, the latent genomic sources of variation in GWAS output may reflect distinct biological pathways influencing for example myelination, synaptogenesis, neuroinflammatory processes, cell division, or cell death, all of which influence many brain IDPs but may act relatively independently. Genomic ICA is therefore based on the premise that the genetic variants influencing the same biological processes will impose more similar associations across thousands of brain IDPs, while SNPs associated with distinct biological processes have different patterns of associations across IDPs. While this form of latent structure of genomic effects has been explored previously in expression data (Kong et al., 2008; Wang et al., 2021), it has not been applied to genome-wide allelic effects on polygenic traits or had its reproducibility tested. We posit that genomic PCA and ICA may furthermore improve upon univariate GWAS output by capturing systematic ‘noise’ (e.g. residual population stratification) that may not have been completely accounted for in the original GWAS quality control. Additionally, genomic ICA may capture genomically distributed components that reflect consistent effects that are more environmentally mediated but have nevertheless been shown to be heritable, such as sociodemographic metrics or lifestyle factors (Abdellaoui et al., 2019; Marees et al., 2021).

Alternative approaches for investigating multi-locus genome-wide associations have been considered to identify hidden patterns in the complex genetic signal of a large GWAS. Multivariate methods, such as latent factor models, including genomic structural equation modelling (SEM) are promising. Genomic SEM tests the fit of *a priori* defined latent factors to the space of multiple phenotypes (Grotzinger et al., 2019) and has been applied successfully to multiple psychiatric (Grotzinger et al., 2019; Marees et al., 2021) and cognitive traits (Warrier et al., 2019), and provided new insights into shared genetic effects across multiple, related traits (Thorp et al., 2020). However, the need to predefine latent factor structures limits the discovery of unhypothesised patterns in the data, and the computational demands of genomic SEM, generally limit the model order to up to ∼20 traits. Alternatively, another new method, Multivariate Omnibus Statistical Test (MOSTest) integrates multiple phenotypes in a single powerful GWAS by combining GWAS test-statistics in a manner akin to meta-analysis, but does not attempt to disentangle the overall SNP-effects across traits into separate mechanisms or inform on which genetic factors modulate different traits in concert (van der Meer et al., 2020). Other recent work has explored the application of PCA to genome-wide variant z-scores among multiple phenotypes to decompose a number of latent genomic factors (Sakaue et al., 2021). Given a set of genetically correlated traits, PCA explains most of the shared variation in as few components as possible. Another recent application used PCA as a data-driven extension of genomic SEM tools, to derive the most salient component (first principal component) from cross-trait genetic correlation matrices, and thus identified patterns of shared genetic variance between multiple MRI traits (Fürtjes et al., 2023). This method by Fürtjes et al. is closest to our proposed methods here. Differences are that it is based on LD-score regression derived genetic correlation matrices, it was only used to extract the first principal component and not to disentangle the signal into multiple (potentially more specific) biological mechanisms, and it was not extended to ICA. However, methods like PCA leverage the shared pleiotropic signal across the traits to derive multivariate genetic components that capture much of the signal very efficiently and likely reduce the influence of noise in the process. Extensions to classical PCA, such as the previously discussed genomic ICA, will improve upon these multivariate methods, and have been underrepresented in the space of genomic analyses.

In the present paper, we present genomic PCA and ICA as novel, fully data-driven methods to decompose large, high-dimensional GWAS summary statistics. This work has evolved from our previous pilot work (Soheili-Nezhad et al., 2021) and is therefore based on the same underlying rationale. In the present study, we present for the first time our complete and final methodological approach of genomic ICA, extended with PCA, along with a systematic evaluation of the robustness under a multitude of different analytic parameters, dimensionality of the output, and other methodological considerations. We test our methods on the GWAS output of 2,240 brain MRI traits from UK Biobank, with large, non-overlapping discovery (n=22,138) and replication (n=11,086) samples (Smith et al., 2021). We determine the reproducibility for multiple versions of our method with varying analytic parameters. We evaluate the output at multiple dimensions (numbers of components). We also apply genomic PCA and ICA to both raw univariate GWAS SNP effect betas, and z-transformed GWAS SNP effect betas. Lastly, we provide a head-to-head comparison of genomic ICA and genomic PCA reproducibility with the reproducibility of corresponding univariate GWAS that was used as input. This work demonstrates the robustness of genomic PCA and ICA to decompose GWAS signal capturing hidden genomic sources of individual differences in variation across thousands of heritable brain traits.

## Methods

### Data

We acquired GWAS summary statistics from the Oxford Brain Imaging Genetics (BIG-40) database. BIG-40 contains the results of over 4000 GWASs that were performed using brain imaging (MRI) derived phenotypes (IDPs) in the UK biobank consortium (UKBB) (Alfaro-Almagro et al., 2018). For this study, two releases of GWAS summary statistics from the UKBB, with 11,086 (11k) and 22,138 (22k) participants respectively, were retrieved (Smith et al., 2021). These samples are non-overlapping and contain an identical set of IDPs, making them well suited for discovery and replication. Due to low heritability of node-to-node functional connectivity metrics (Elliott et al., 2018) we excluded these from the analysis, leaving 2240 IDPs for further analysis. Notably, the amplitudes of functional signal fluctuations and six ICA-derived ‘global’ measures of functional connectivity were included, as they were shown to be heritable (Elliott et al., 2018; Smith et al., 2021). Other IDPs included metrics from classes of T1-weighted MRI, diffusion MRI, susceptibility-weighted imaging (SWI), fluid-attenuated inversion recovery (FLAIR), task MRI and quality control (QC) procedures. For details on the GWAS pipeline of the UKBB please refer to the main publications (Elliott et al., 2018; Smith et al., 2021) and the website including these UKBB releases (https://open.win.ox.ac.uk/ukbiobank/big40).

### Clumping

We applied genome-wide SNP clumping to reduce local SNP dependencies stemming from linkage disequilibrium (LD). First, we pruned all available SNPs with a threshold of r^2^<0.3. Then, we considered the smallest p-value of each SNP across the GWASs of 2,240 brain IDPs for clumping. A genomic window size of one mega base, a lead variant p-value threshold of 10^-5^ and a LD-threshold of r^2^>0.1 were used as clumping parameters. LD was estimated in a random subsample of 10,077 Caucasian UK biobank participants (data field number 22006). This procedure reduced the total number of genome-wide SNPs from n=17,103,079 to n=157,893. Thereby, we minimise the impact of LD on local SNP-to-SNP correlations while keeping brain-related lead variants within each LD block. This clumping procedure is in line with the consensus in the field, whereby we deem within-chromosome lead-SNPs associated with genetic loci as independent at r^2^<0.1 (Watanabe et al., 2017). To make the methodology more sensitive to genetic effects of common neurological and psychiatric disorders for future downstream analyses, we fortified the n=157,893 clumped SNPs with additional lead SNPs associated with Attention Deficit/Hyperactivity Disorder (ADHD) and Alzheimer’s Disease (AD), thereby introducing an additional n= 7,471 SNPs into the analysis. This increased the number of SNPs from n=157,893 to n=165,364.

### Genomic ICA and genomic PCA

We concatenated all GWAS regression values, representing the genome-wide SNP effect sizes, across all IDPs, generating an m×n brain-wide genome-wide matrix of imaging IDPs (m=2,240) and genetic variants (n=165,364 clumped SNPs). Subsequent multivariate decomposition was applied to two versions of the m×n brain-wide genome-wide matrix: the raw GWAS betas, and the z-transformed GWAS betas quantifying SNP effect sizes standardised for their own standard errors. We decomposed the brain-wide genome-wide matrices of SNP effect sizes using v3.15 of Multivariate Exploratory Linear Optimized Decomposition into Independent Components (MELODIC), which is a probabilistic ICA algorithm maximising non-Gaussianity in the reconstructed independent sources (Beckmann & Smith, 2004). In the standard pipeline, MELODIC starts by applying probabilistic PCA to the data, thereby decomposing the data into a predefined number of principal components. Then, the algorithm rotates these principal components to optimise a measure of independence (i.e. non-Gaussianity), thus producing the same number of independent components. Each extracted component consists of a latent genomic factor of SNP-loadings in the SNP dimension, and a vector of IDP-loadings in the MRI dimension. This feature allows us to determine the association of individual IDPs with the corresponding hidden genomic factor. The m×n brain-wide genome-wide matrices were decomposed by MELODIC along the SNP dimension into a maximum of 200 principal and independent components of SNP effect sizes. For further analyses dimension 50 was chosen as maximum dimension via visual assessment of the scree plots (Figures 2 and S2) generated from dimension 200, which indicated that dimension 50 provides a good balance between explained variance and model complexity (Figure 2, ∼61% SNP effect variance explained in 33k sample). Inter-sample reproducibility was calculated for 5, 10, 25, and 50 dimensions in the discovery (22k) and replication (11k) samples for both z-transformed and raw beta input matrices. We disabled the global signal removal option of MELODIC since SNP effect size distribution is already centred on zero under the null assumption, given the random direction of effects dependent on the effect allele. Furthermore, we disabled SNP variance normalisation across IDPs to preserve the magnitude of allelic effects in the genomic components which contain biological information. This is in contrast to MELODIC applications to fMRI data, where the inherently relative nature of the data warrants the variance normalisation step (Beckmann & Smith, 2004).

### Reproducibility testing

To determine reproducibility of principal and independent genomic sources in independent samples across component pairs we focused on two measures: First, to assess the non-sparse, global genomic signals of all variants contributing to each component, we calculated Pearson’s correlation coefficients of SNP-wise loadings. The statistical associations were corrected for multiple comparisons by Bonferroni correction for the number for all unique comparisons, here N^2^, where N is the number of components per decomposition.

Secondly, to focus only on the sparse part of the genomic sources (the tails of the distribution), SNPs that strongly contribute to each component’s multivariate effect relative to the loading distribution, while removing possibly noisy low end of the loadings, we binarized and thresholded the component loadings at values >1. To determine statistical significance of the degree of overlap between each component from the discovery sample with each component of the replication sample we performed a Fisher’s Exact Test, adjusting for multiple comparisons by the number of contingency tables generated. Fisher’s exact test is exact under the assumption that lead-SNPs with r^2^<0.1 are statistically independent, such that the expected degree of overlap under the null can be calculated. Given some degree of LD was still present (at r^2^<0.1), we also derived the number of effectively independent SNPs (Galwey, 2009) to determine if our analysis should be adjusted to account for this slight remaining dependence between lead-SNPs. The method and the outcome of this analysis are described in the supplement. From this supplementary analysis, we concluded that the difference between the number of effective (independent) SNPs given our LD threshold, and the actual number of SNPs, was negligible.

### Reproducibility of univariate GWAS

We determined reproducibility of raw, univariate GWAS SNP effect sizes and z-transformed, univariate SNP effect sizes separately to provide a benchmark for the decompositions of both versions of input data. We computed the Pearson’s correlation coefficient (*r_SNP_*) of variant effect sizes across independent samples. Maximum r^2^_SNP_ is theoretically determined by additive SNP heritability (*h^2^_SNP_)* of each trait, and provides a benchmark for assessing the reproducibility of PCA and ICA genomic components.

### Visualisation of IDP clusters of independent genomic components

The decomposition of large MRI-GWAS data with MELODIC yields a set of IC loadings that quantify the covariation in IDP-space in correspondence to the covariation of genetic effects in the SNP-space. The IDP-space of N=2,240 IDPs can be divided into general classes of MRI modalities, namely cortical surface area, cortical thickness, diffusion MRI derived metrics using tract-based spatial statistics (TBSS) and probabilistic tractography approaches, grey matter volume assessed via FMRIB’s Automated Segmentation Tool (FAST), subcortical ROI volume assessed using FMRIB’s Integrated Registration and Segmentation Tool (FIRST), ROI volume across multiple atlases assessed using Freesurfer, task-based functional MRI, and resting-state functional MRI. To visualise the clustering in IDP-space driven by genetic effects in SNP-space, we embedded the IDP-loadings into a two-dimensional space using t-distributed stochastic neighbour embedding (t-SNE) (Maaten & Hinton, 2008). To maximise the clustering potential of t-SNE the visualisation was performed on the decomposition of the combined sample of 11k and 22k UKB samples.

## Results

### Raw and z-transformed univariate GWAS

Reproducibility of z-transformed, univariate GWAS betas across samples ranged from r_max_=0.28 to r_min_=0.005 (r_mean_=0.11). Highest reproducibility was found in phenotypes derived from large white matter tracts derived from *dMRI*, followed by IDPs in global *cortical volume*, *thickness*, and *surface area. Cortical surface area, intensity measures* and *FAST-derived* measures showed lower reproducibility than *cortical thickness* and *volume.* Lowest reproducibility was found for *SWI* and *fMRI* derived IDPs. Reproducibility of all IDPs is shown in Figure 1. Reproducibility of raw, univariate GWAS beta-values in terms of Pearson’s correlation coefficient, ranged from r_max_=0.25 to r_min_=0.003 (r_mean_=0.09) and was lower than z-transformed univariate GWAS reproducibility. Highest reproducibility was found for similar IDPs as for the z-transformed decomposition and depicted in figure S1.

**Figure 1:**
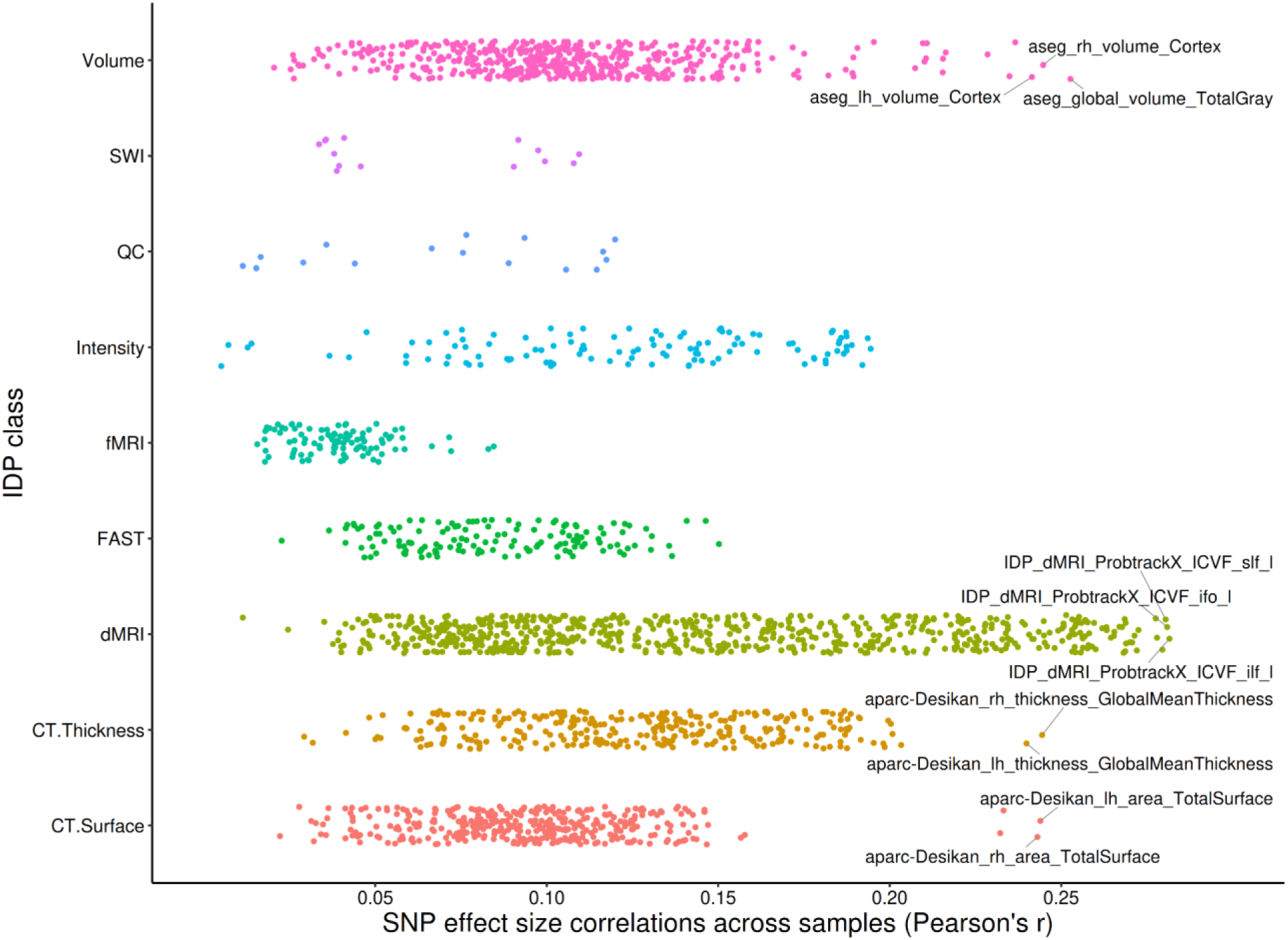
Reproducibility of the z-transformed, univariate GWAS SNP effect sizes (n=165,364 clumped variants) across independent samples.

### Principal genomic components

The first five PCs derived from z-transformed GWAS captured 31.9% of the variance across SNP effect sizes, while decomposing into 200 PCs increased the variance explained to 79.6% (Figure 2). A nearly identical pattern was found for the variance captured by the non-transformed raw GWAS betas (Figure S2). Inter-sample reproducibility of PCs at dimension 5 was high, ranging from |r_max_|=0.33 (*p_adj_*=<10^-308^) to |r_min_|=0.18 (*p_adj_*=<10^-308^), with decreasing reproducibility at higher dimensions (figure 3A & B; table S1). Notably, the first PC was more reproducible than the maximum reproducibility across all 2,240 univariate z-transformed GWAS outputs (figure 3B). The first ten PCs derived from z-transformed GWAS all showed higher reproducibility than mean reproducibility of raw z-transformed GWAS (figure 3A & B). Subsequent PCs #11-#50 successively explained less variance, and were also less reproducible across independent samples (figure 3B). The reproducibility of PCs derived from raw GWAS data overall showed a similar pattern but was generally lower than the PCs derived from z-transformed GWAS data (table S1).

**Figure 2:**
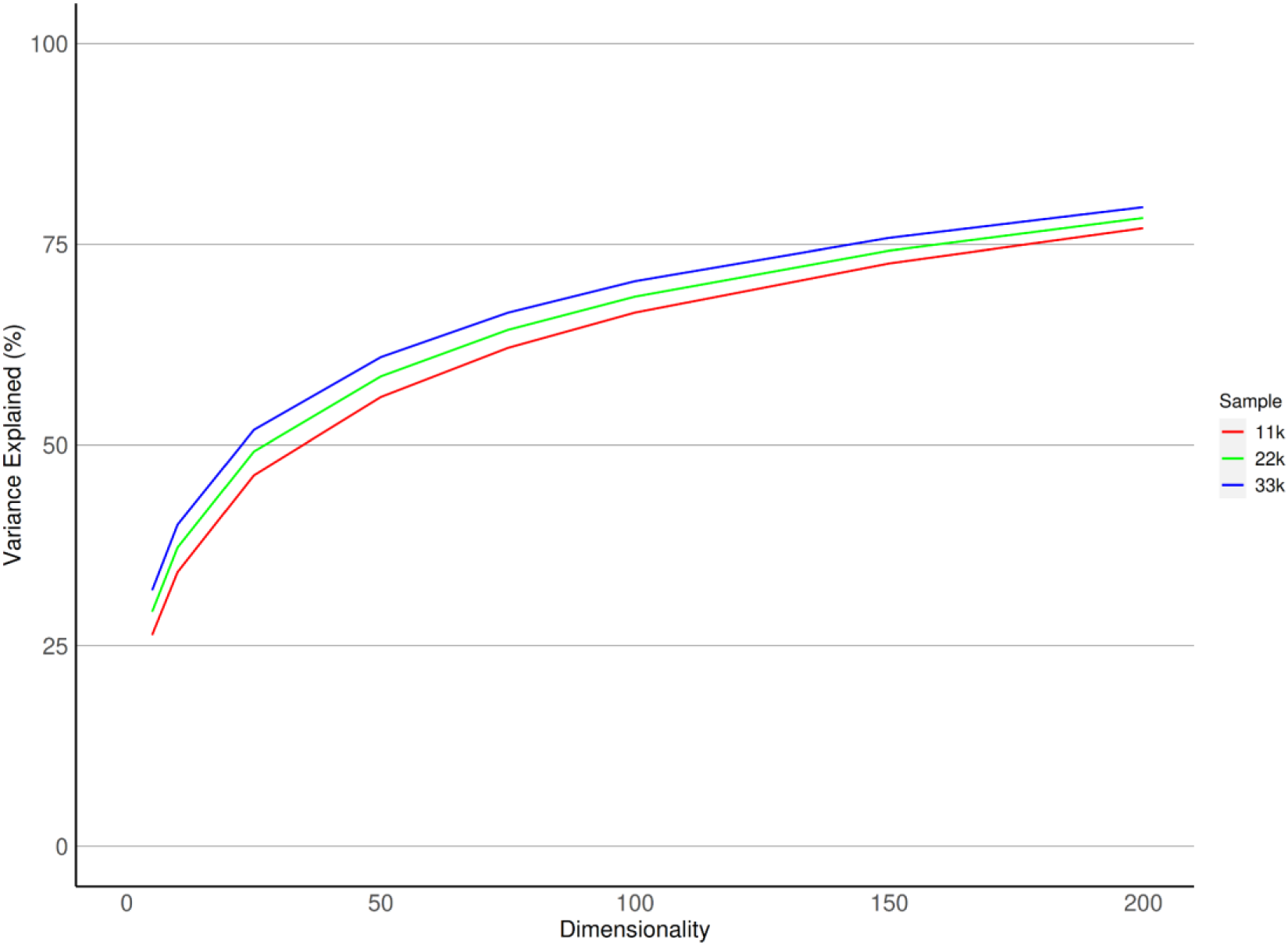
Variance explained by genomic components derived from z-transformed, univariate GWAS SNP effect sizes at PCA dimensions 5, 10, 25, 50, 100, 150, and 200.

**Figure 3:**
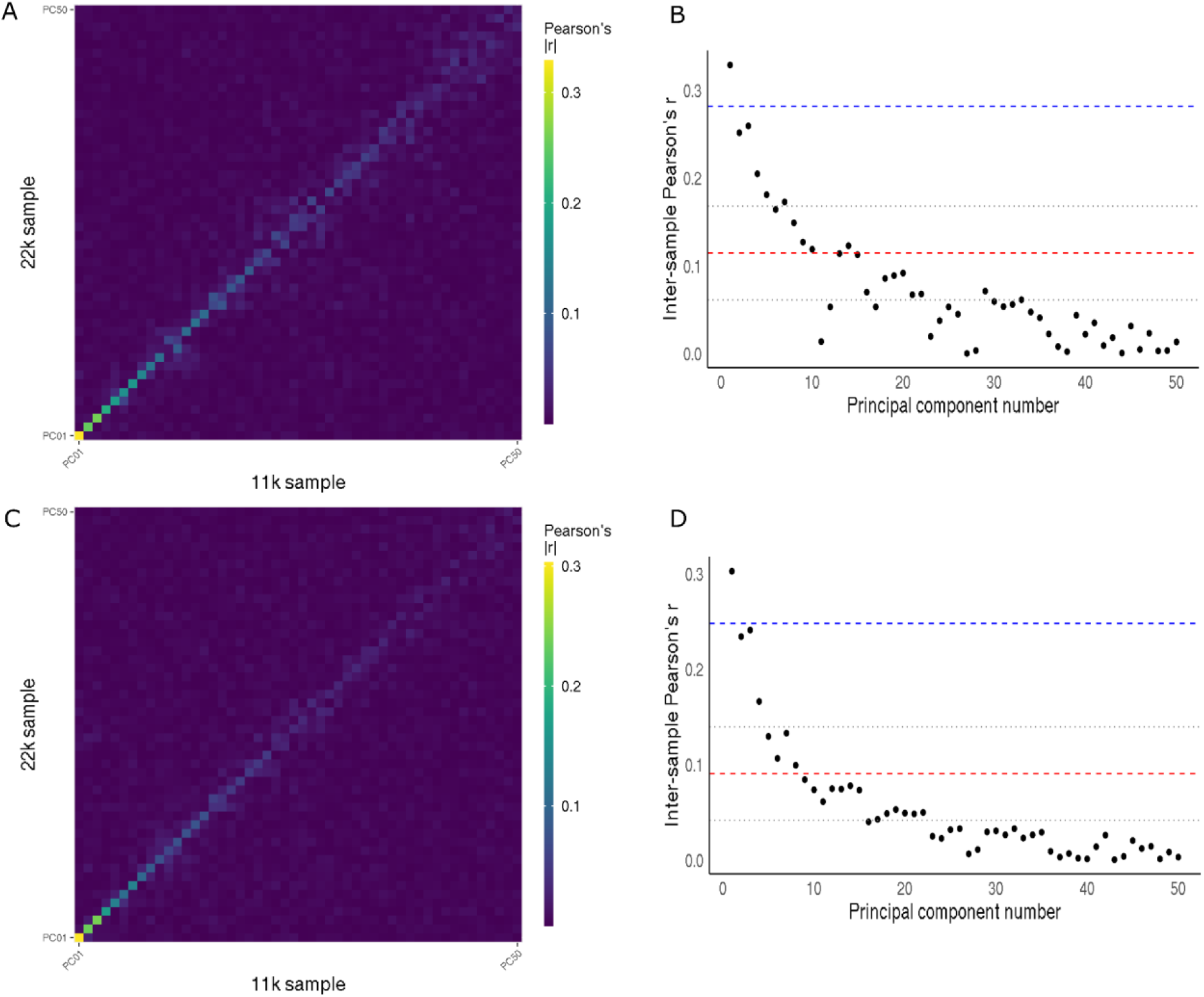
Inter-sample reproducibility of principal genomic components derived at dimension 50, from z-transformed (top row) and raw (bottom row), univariate GWAS SNP effects, displayed as the Pearson correlation coefficient. B and D show the respective reproducibility per independent component as a scatterplot, with the Pearson correlation coefficient on the y-axis. The red dashed line denotes the mean of raw, univariate GWAS reproducibility, with the grey, dotted lines indicating one standard deviation around the mean. The blue dashed line indicates the maximum reproducibility of z-transformed, univariate GWAS betas.

### Independent genomic components

ICs derived from z-transformed, univariate GWAS likewise showed highest reproducibility at dimension 5 (|r_max_|=0.25, *p_adj_*=<10^-308^; |r_min_|=0.15, *p_adj_*=<10^-308^; |r_mean_|=0.20; figure S3A & B). Reproducibility dropped with increasing dimensionality (figure S4 & S5). Improved reproducibility compared to univariate GWAS was found up to dimension 10 (|r_max_|=0.23; |r_min_|=0.12; |r_mean_|=0.16; figure 4A & B). At dimension 10 we found that all components from the discovery sample correlated with either one or multiple replicated components with high statistical significance (0.23 > |r| > 0.10; p_all_=<10^-308^; table S2). ICs derived from z-transformed GWAS data were more reproducible than ICs derived from raw GWAS data (table S2). While the maximum reproducibility among the 10 ICs was lower than the maximum univariate reproducibility among 2,240 IDPs (|r_IC2_|=0.23 vs r=0.28), all 10 ICs exceeded mean univariate reproducibility (figure 4B). After binarizing, the top SNPs of six of the ten independent components of the discovery sample replicated significantly (*Fisher’s p*_all_=<0.003; table S2). The most strongly correlated, IC 1 from the discovery sample replicated as IC 2 in the replication sample (*Fisher’s p_adj_*=5.5*10^-66^). Reproducibility statistics of genomic independent components at dimensions 5, 25 and 50 and a comparison with respective, univariate GWAS reproducibility are shown in the supplement (figure S3 – S5).

**Figure 4:**
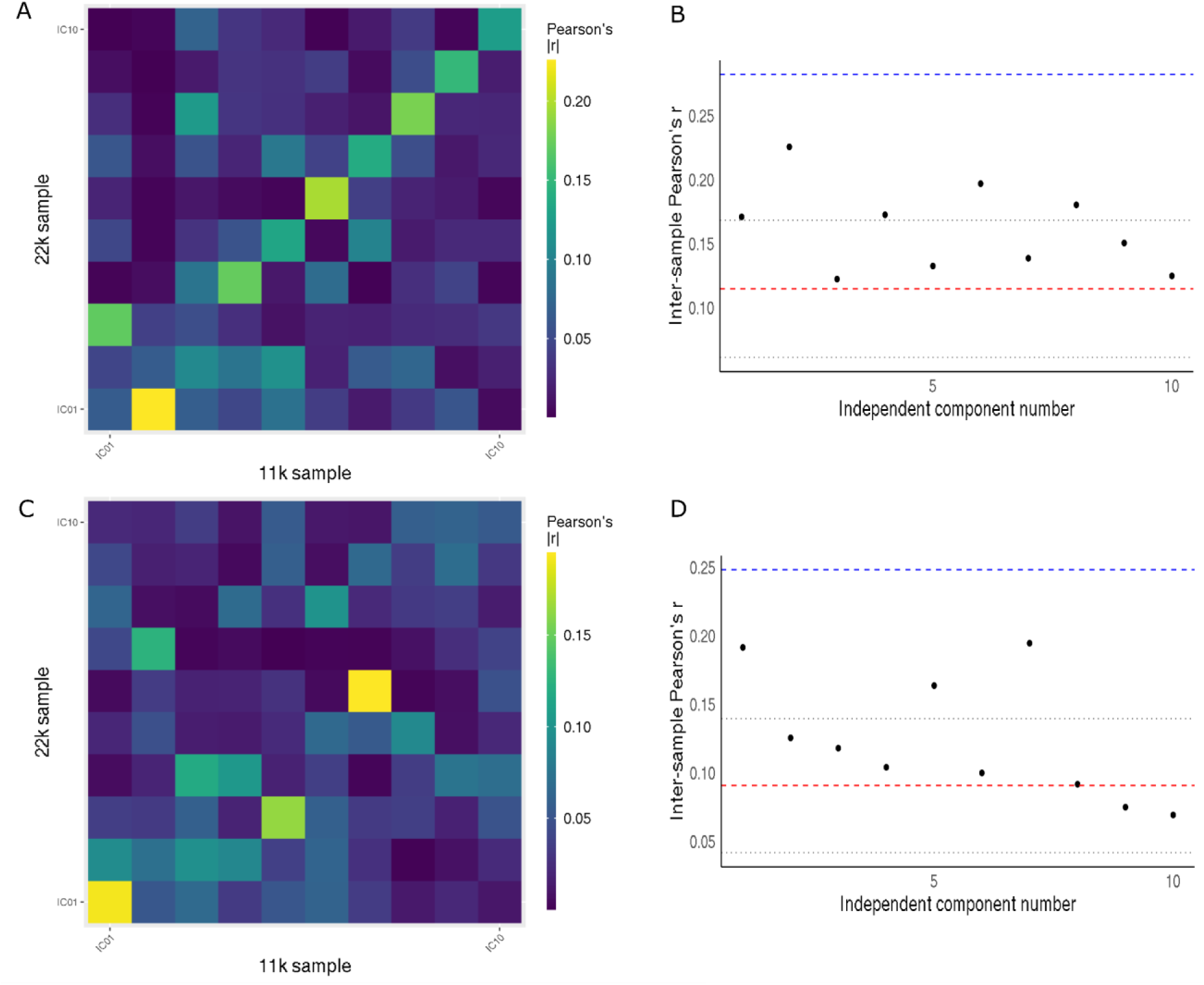
Inter-sample reproducibility of independent genomic components derived at dimension 10, from z-transformed univariate GWAS SNP effects (A), and raw univariate GWAS SNP effects (C) displayed as the Pearson correlation coefficient. B shows the reproducibility per independent component derived from z-transformed univariate GWAS as a scatterplot, with the Pearson correlation coefficient on the y-axis. D shows the reproducibility of raw univariate GWAS in the same manner. The red dashed line denotes the mean reproducibility of the respective univariate GWAS, with the grey, dotted lines indicating one standard deviation around the mean. The blue dashed line indicates the maximum reproducibility of the respective univariate GWAS betas.

Independent components derived from raw, univariate GWAS followed a similar pattern as z-transformed decomposition (figure 4C & D; figure S3 - S5). At dimension 10 eight components exceeded mean reproducibility of univariate GWAS (|r_max_|=0.19, |r_min_|=0.07, |r_mean_|=0.12, figure 4D). Components from the discovery sample correlated with one or multiple components in the replication sample with high statistical significance (0.19 > |r| > 0.07; p_all_=<1.88^-157^, table S2). The distribution of component loadings from raw GWAS decompositions were highly kurtotic, which made the investigation of overlap of the component tails unsuitable. Reproducibility of dimensions 5, 25 and 50 and a comparison with raw, univariate GWAS SNP effect reproducibility are shown in the supplement (figure S3 - S5).

### IDP clustering of independent genomic components

The reproducibility analysis showed that the decomposition of z-transformed, univariate GWAS at dimension 10 yielded the best balance between model complexity and inter-sample reproducibility among the decomposition parameters tested. The t-SNE analysis of the corresponding IDP loadings clearly showed clustering of IDP loadings along the boundaries of larger IDP groups in MRI modalities (Figure 5). These results indicate distinct associations of concerted genetic effects on traits from different modalities and similar genetic effects within imaging modalities (e.g. cortical thickness and cortical surface area vs. dMRI measures). In some cases different methods used to derive metrics related to similar modalities resulted in the splitting into different clusters, such as with T1-weighted images of cortical thickness and surface area, SWI and MRI Intensity IDPs. Different methods in diffusion MRI did not follow this trend, and probabilistic tractography derived IDPs and TBSS derived IDPs showed clustering according to similar genetic associations.

**Figure 5:**
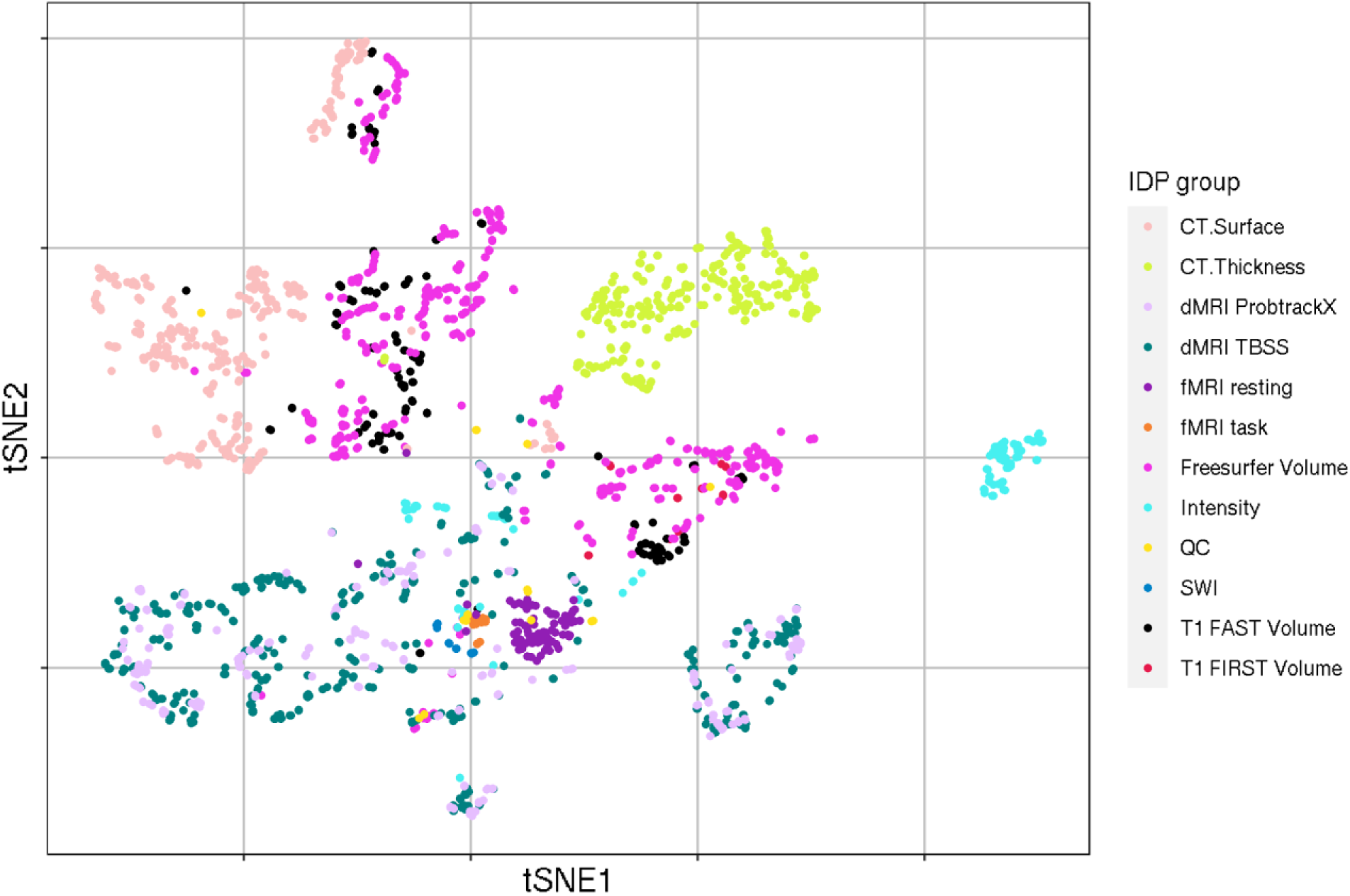
t-SNE based visualisation of IDP loadings, derived from the decomposition into 10 genomic ICs of z-transformed univariate GWAS of the combined 11k and 22k samples. This plot shows the clustering of IDP loadings across all dimension 10 genomic components, thereby showing the emergence of distinct IDP groups associated with the covariation of specific sets of genetic effects.

## Discussion

GWAS analyses of the past decades have enhanced our understanding of common genetic variants influencing imaging derived brain phenotypes. The complexity of gene functions and their many interactions with proteins, cells, and environment obscure the biological pathways underlying genome-wide associations with brain measurements. Further, pleiotropic and polygenic effects, paired with small effect sizes of genetic variants, limit the mechanistic interpretability of GWAS data. Genomic PCA and ICA take advantage of pleiotropy and polygenicity through the method’s assumption that the GWAS effect sizes across genetically correlated traits are a linearly mixed signal containing structured and gaussian noise. This makes these methods well suited to uncover hidden structure within large GWAS data on over 2,000 IDPs. The mathematical framework of genomic PCA and ICA provides a way to disentangle the complex web of genetic effects by grouping SNP effects into genetically covarying clusters. This could enhance our understanding of how genes act by presenting novel gene-sets based on genomic ICA and PCA for downstream analysis.

We decomposed the present GWAS data of over 2000 IDPs, without *a priori* assumptions, into a smaller set of multivariate principal and independent components. We tested the reproducibility of genomic components across independent samples to assess robustness of the proposed method. We identified reproducible genomic components, in both the raw, univariate GWAS SNP effects and the z-transformed univariate GWAS SNP effects. We found that genomic components derived from both data sets show moderate reproducibility across independent samples, substantially improving upon mean reproducibility of univariate GWAS. For both raw and z-transformed betas, PCA captured most of the variance within the first three components, with the following components capturing less variance. Reproducibility of the first independent genomic components was lower than of the principal components, but at dimension 10 all ICs also showed improved reproducibility across all IDPs compared to mean reproducibility of respective GWAS data up to dimension 10. IDPs from distinct MRI modalities show clear clustering patterns, indicating distinct genetic effects acting on MRI-modality and tissue specific brain IDPs (figure 5).

Reproducibility of genomic PCs and ICs derived from z-transformed data was higher than those of components derived from raw beta-values. This is likely because the z-transform accounts for the variable standard error around the SNP-betas making them less sensitive to noisier SNP estimates with large standard-errors, which would be especially the case for low-MAF SNPs. With increasing discovery sample sizes for GWAS the standard errors shrink, ultimately leading to convergence of raw and z-transformed decompositions to identical results. Increases in discovery sample sizes will also enable decompositions into more and more reliable independent sources of genomic signal, thereby increasing the sensitivity of genomic PCA and ICA to uncover more refined and likely more specific clusters of genetic effects on sets of brain features.

The ultimate goal of the application of genomic PCA and ICA is to uncover hidden structure in genetic effects on brain IDPs, to enhance our understanding of the biological mechanisms driving brain development and the emergence of mental illnesses. The hypothesis-free, data-driven approach is inherent to genomic PCA and ICA, thereby not relying on predefined gene-sets or molecular pathways to determine effects on IDPs. The components presented here are a hidden representation of the genetic architecture subserving over 2000 brain IDPs. The comparison of genomic PCA and ICA showed that they are similarly reproducible across independent samples, with the exception of the first three (most salient) PCs with the highest reproducibility. While genomic ICA yielded components that were less reproducible than PCA, they were still a marked improvement on univariate GWAS considering we decomposed across all SNPs and IDPs simultaneously. The drop in reproducibility after dimension 10 is likely due to the distribution of genomic, independent signals across increasingly many components, as dimensionality increases. This trend can already be observed at dimension 10, where some components from the discovery sample correlate significantly with multiple replication components. Theoretically, PCA extracts orthogonal axes of maximum variance from signal, thus Genomic PCs likely represent a convoluted mixture of many underlying biological signals across IDPs. In contrast, ICA separates hidden, independent sources of variation from mixed signals containing Gaussian noise. This allowed us to conclude that our present findings are consistent with the theoretical framework. Univariate GWAS reproducibility was highest for global measures of cortical morphology and large tracts of white matter architecture. Specifically, highest univariate reproducibility of both “raw” and z-transformed GWAS SNP effects were found in measures of global *cortical volume*, *cortical thickness*, and metrics of large white matter strands. These findings echo earlier heritability estimates of white matter fractional anisotropy that found larger *h^2^*for large white matter structures (e.g. superior longitudinal fasciculus, *h^2^*∼0.87) compared to smaller structures (e.g. fornix, *h^2^*∼0.53) (Kochunov et al., 2015). The GWAS signal representing the global features may thus be influenced by “signal averaging” across subregions, leading to relatively less noise and increased phenotypic reproducibility. Genomic ICA and PCA are thus theoretically well suited to further enhance signal to noise in a similar but non-additive way by accumulating shared signal from IDPs affected by similar genetic effects in the same component. This suggests that genomic ICA and PCA may be effective in separating biological signals from structured and unstructured (gaussian) noise, as is also the case in classic neuroimaging applications (Pruim et al., 2015).

ICA is a method that decomposes large data without a priori assumptions. Theoretically, there is no limitation for which data to use this method on. Therefore it can be used on fMRI data (Beckmann & Smith, 2004) and, here, on genomics data to separate out hidden independent sources of relevant signal. Furthermore, the more data is available for ICA to parse the better the potential signal in the data can be separated from similarly present structured noise. As such ICA thrives on more potential signals hidden in data, as long as the structured noise permeates the same data. This makes a case for including non-MRI based GWAS data in genomic PCA and ICA decompositions as it would further improve discoverability of more fine-grained sources of genetic variation from large GWAS. The flexibility of PCA and ICA could also be applied to epigenome-wide and transcriptome-wide data, thereby deepening our understanding of environmental effects on genetic expression patterns. Additionally, future research could explore other, powerful means of decomposition that are sensitive to weightings of included modalities by decomposing across data domains using, for example, linked ICA (Groves et al., 2011) or SuperBigFlica (Gong et al., 2022). Another avenue that we currently pursue is the computation of individualised component scores, following the polygenic risk score framework, to stratify existing cohorts for normative modelling and personalised medicine. We anticipate that with increased sample sizes of discovery neuroimaging GWAS data, the components will become more reliable and interpretable by capturing more fine-grained, covarying genetic effects. Investigations into the alignment of component IDPs with gene-sets based on cell specific gene expression, molecular pathways, or brain homeostasis will reveal the potential of our methods.

## Limitations

Association of GWAS SNP effects decomposed with genomic ICA may still influence one another on any level of the causal chain from DNA molecule to fully developed brain IDP, and thereby may conceal effects. This touches on the question of what noise means in the context of genetic data. While genomic PCA and ICA are well suited to extract structured noise from data and capture it in individual components, we can not be certain how this structured “noise” affects brain development on any level of biological, causally related mechanisms. Some components might capture structured noise insufficiently captured by quality control protocols, such as population stratification, assortative mating, or cryptic relatedness. Other components may capture a mixture of genetic effects related to “house-keeping” mechanisms that affect brain-wide mechanisms which influence diverse tissue types and properties. This warrants further analyses of reproducible components using for example gene-set enrichment and other bioinformatics tools. Even though we decompose a large matrix containing ∼2000 brain IDPs, the inclusion of non-MRI based data may further improve the outcome of genomic ICA by providing the method with more data to parse, as discussed above. This is different from the approach used by Fürtjes et al., which is closest to our proposed methods here, where only structural MRI derived volumetric IDPs were used (Fürtjes et al., 2023). While the application of PCA is common between their and our work, this constraint in the IDP space might have limited their discoverability of distinct genetic components that are shown to map to distinct modality clusters (figure 5). This shows the potential of using data-driven, hypothesis free methods on large GWAS-MRI data to uncover hidden structure across IDPs. The inclusion of non-MRI based data in addition to the ∼2000 brain IDPs may reconstruct yet more sources of genomic variation related to behavioural measures, disease biomarkers and psychiatric conditions. Further, the present analysis using GWAS data is the Western European-centric ancestry of the UK Biobank sample. These components may be more or less sensitive to genetic effects specific to genetic ancestries and socio-cultural influences. We recognize the efforts to expand GWAS to international cohorts which will provide more data for these genetic analyses, and we eagerly await the availability of these data to be included into the genomic PCA and ICA framework.

## Conclusion

We introduce genomic PCA and ICA as a novel method to decompose large MRI-GWAS summary statistics to efficiently and reproducibly extract latent genomic components that affect brain structure and function as measured by MRI. We derived principal and independent genomic sources from a large set of GWAS statistics, containing genetic associations with 2240 brain IDPs. To thoroughly test the efficacy of the method we decomposed both raw and z-transformed, univariate GWAS SNP effects at multiple component dimensions. Genomic PCA and ICA showed improved inter-sample reproducibility for both decomposition inputs, compared to respective, univariate GWAS SNP effects. Genetic effects captured across components showed clear clustering according to specific MRI modalities and brain features. This makes genomic ICA a promising method to consistently and effectively decompose noisy, brain-related genome-wide association data into more reproducible and more interpretable genomic components capturing covariation of genetic effects across IDPs.

## Acknowledgements

This work was supported by the Horizon Europe grant (No. R0006682) for the FAMILY consortium (http://family-project.eu), funded by the European Union, the Swiss State Secretariat for Education, Research and Innovation (SERI) and the UK Research and Innovation (UKRI) under the UK government’s Horizon Europe funding guarantee. CFB gratefully acknowledges funding from the Wellcome Trust Collaborative Award in Science 215573/Z/19/Z, and the Netherlands Organization for Scientific Research Vici Grant No. 17854 and NWO-CAS Grant No. 012-200-013.

