## Supplemental Material for "Principal and Independent Genomic Components of Brain Structure and Function"

### Additional Analyses

#### Estimation of effective tests

In the present study we reduced the number of SNPs (n=17,103,079) via clumping and pruning to n=157,893, then enriched the sample with another n= 7,471 SNPs related to Alzheimer's Disease and ADHD. This was based on pruning with subsequent clumping, yielding lead variants with LD of $r^2<0.1$. However, the clumped and enriched variants could still be in LD with each other. To account for this we chose to test the number of effective tests conservatively, to estimate if another round of clumping would be necessary to account for non-independence between SNPs.

The number of effective tests (*M*eff) was estimated based on chromosome-specific correlation matrices to account for full SNP independence, in terms of full linkage equilibrium, between chromosomes, which arises definitionally from the concept of LD. The number of effective tests was estimated following the mathematical framework of Galwey (Galwey, 2009), where we calculated the variance of eigenvalues of SNP-correlation matrices per chromosome, and derived the number of effective tests. This method was chosen above another proposed method that requires the manual setting of the eigenvalue (Gao et al., 2008). Given that we aimed to estimate *M*eff in a data-driven way, we decided to use the Galwey method.

Specifically we *M*eff is defined as

$$M_{\text{eff}} = \frac{\left(\sum_{i=1}^{M} \sqrt{\lambda_i}\right)^2}{\sum_{i=1}^{M} \lambda_i}$$

where *M* is the number of tests (or elements in the matrix provided), and lambda denotes the i-th eigenvalues of the matrix given (Galwey, 2009).

To be conservative in light of the inclusion of ~7.000 SNPs post clumping, we assumed correlation matrices, where the diagonal is 1, accounting for full LD of the same SNPs, and with $r^2=0.1$ between all remaining SNP pairings. Another consideration was to use an autoregression algorithm to simulate LD between included SNPs. This idea was rejected however, as extracting the actual LD from the 10,077 UKBB individuals would have been less work and computationally more efficient. Then, we calculated the number of effective tests per chromosome, summed the numbers of effective tests per chromosome to arrive at the full number of effective tests for all n=165,364 SNPs. We found that the total number of effectively independent tests, calculated as specified above, was *M*eff=149,919. This reduced the number of tests by n=15,427. Based on this, and the small p-values we observed when calculating Fisher's exact test on the number of SNPs, we determined that the effect was negligible and another round of clumping was unwarranted at this stage.

### Figures

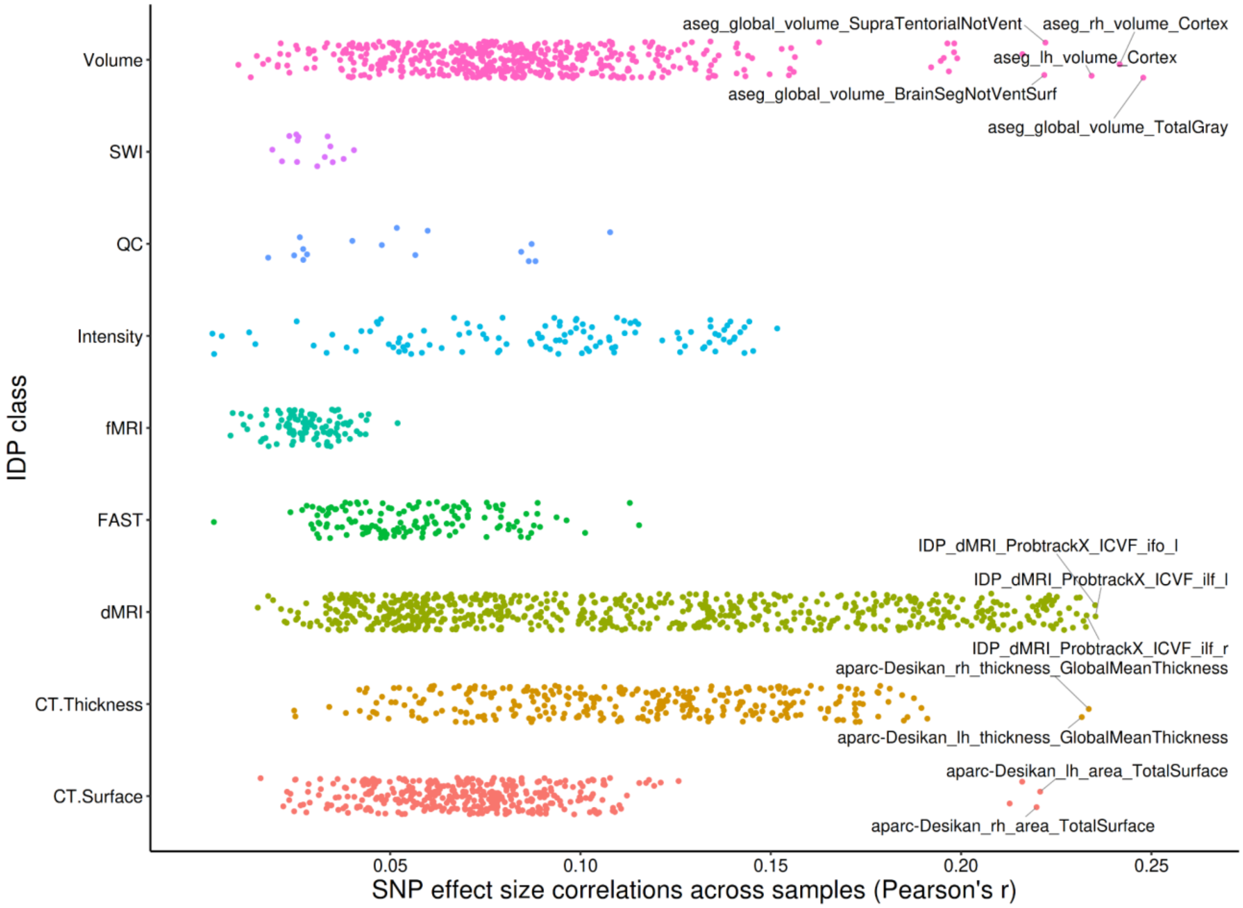


**Figure S1**: Reproducibility of the "raw", univariate GWAS SNP effect sizes (n=165364 clumped variants) across independent samples.

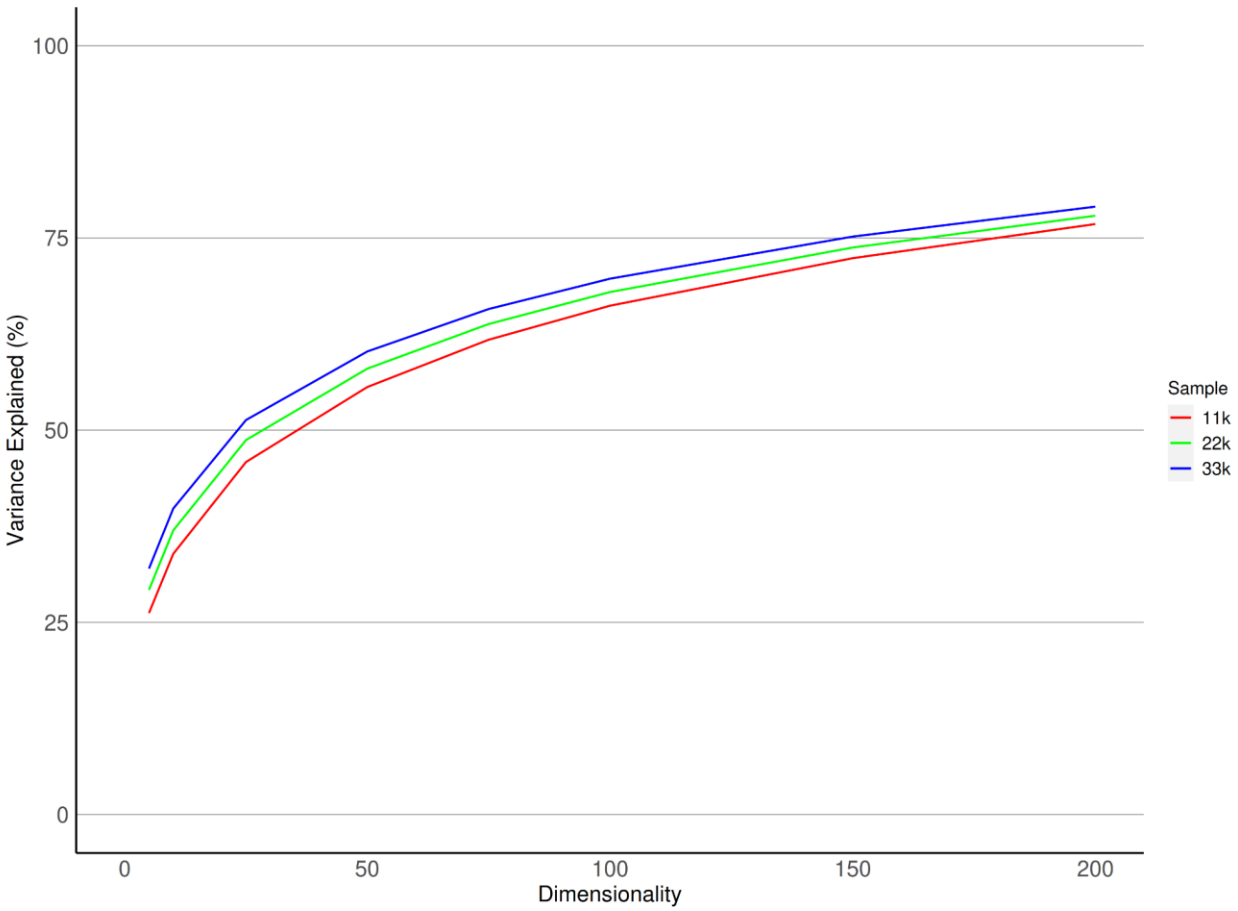


**Figure S2**: Variance explained by genomic components derived from raw, univariate GWAS SNP effect sizes at ICA dimensions 5, 10, 25, 50, 100, 150, and 200.

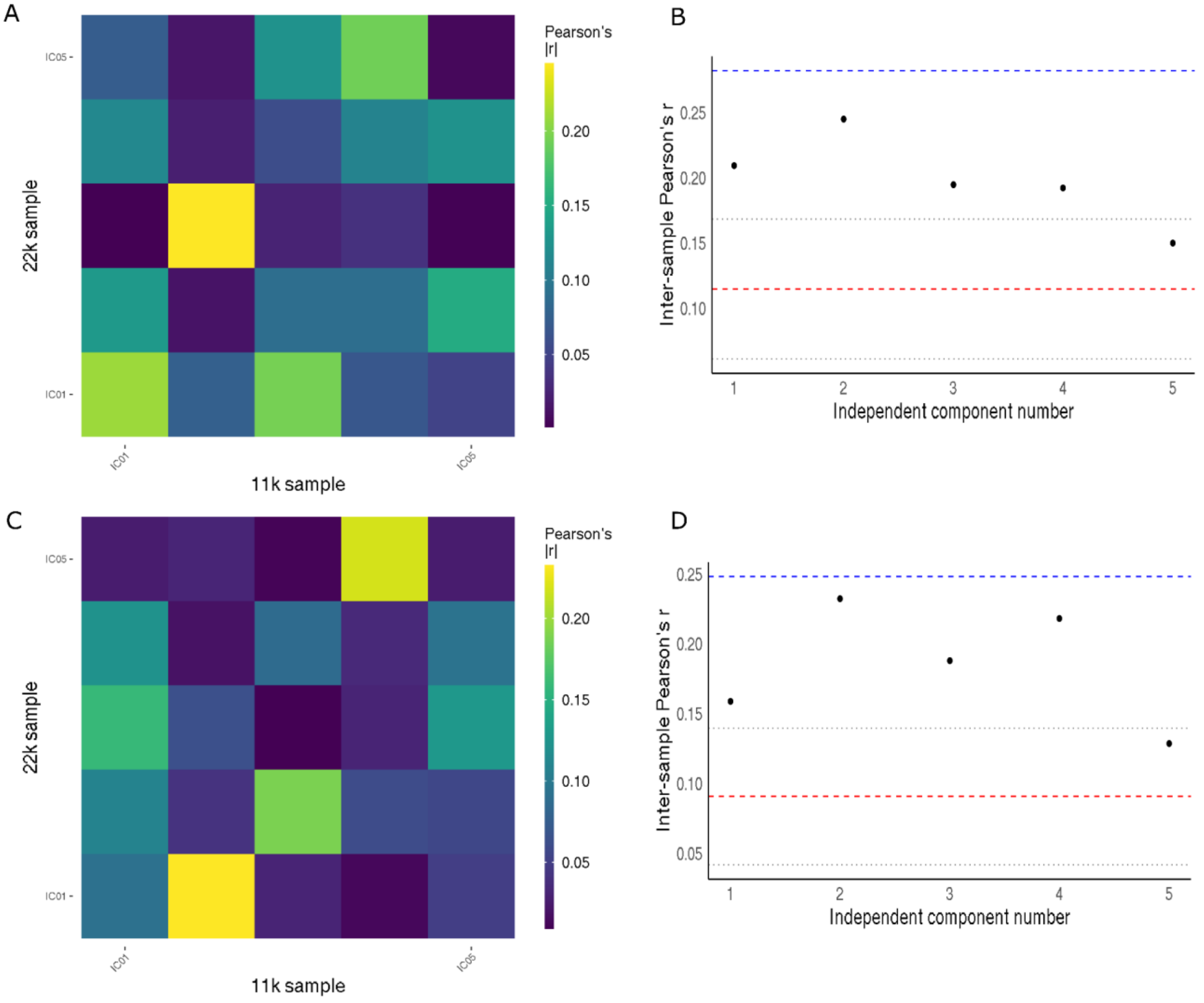


**Figure S3**: Inter-sample reproducibility of independent genomic components derived at dimension 5, from z-transformed (top row) and raw (bottom row), univariate GWAS SNP effects, displayed as the Pearson correlation coefficient in A and C. B and D show the respective reproducibility per independent component as a scatterplot, with the Pearson correlation coefficient on the y-axis. The red dashed line denotes the mean of raw, univariate GWAS reproducibility, with the gray, dotted lines indicating one standard deviation around the mean. The blue dashed line indicates the maximum reproducibility of z-transformed, univariate GWAS betas.

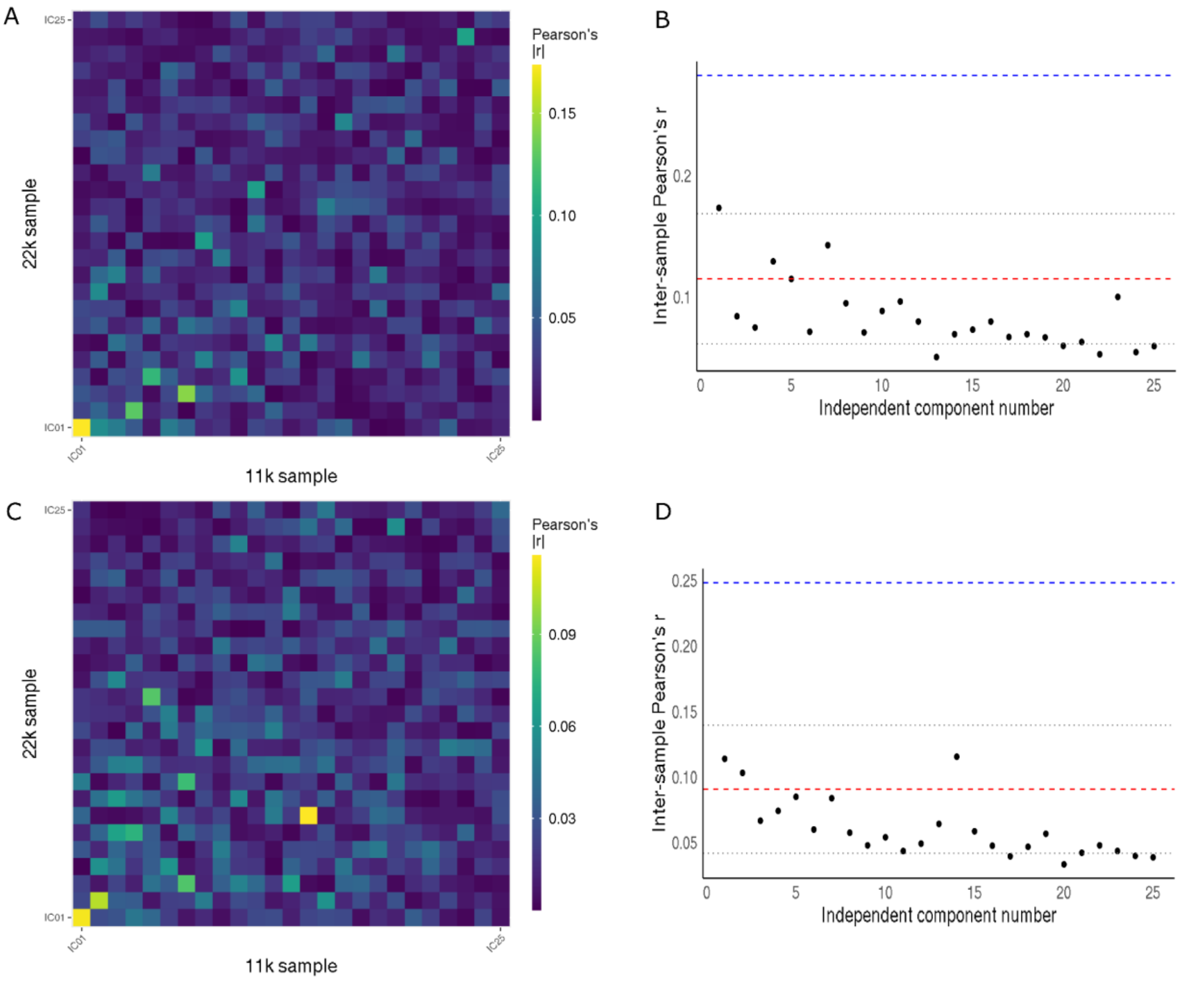


**Figure S4**: Inter-sample reproducibility of independent genomic components derived at dimension 25, from z-transformed (top row) and raw (bottom row), univariate GWAS SNP effects, displayed as the Pearson correlation coefficient in A and C. B and D show the respective reproducibility per independent component as a scatterplot, with the Pearson correlation coefficient on the y-axis. The red dashed line denotes the mean of raw, univariate GWAS reproducibility, with the gray, dotted lines indicating one standard deviation around the mean. The blue dashed line indicates the maximum reproducibility of z-transformed, univariate GWAS betas.

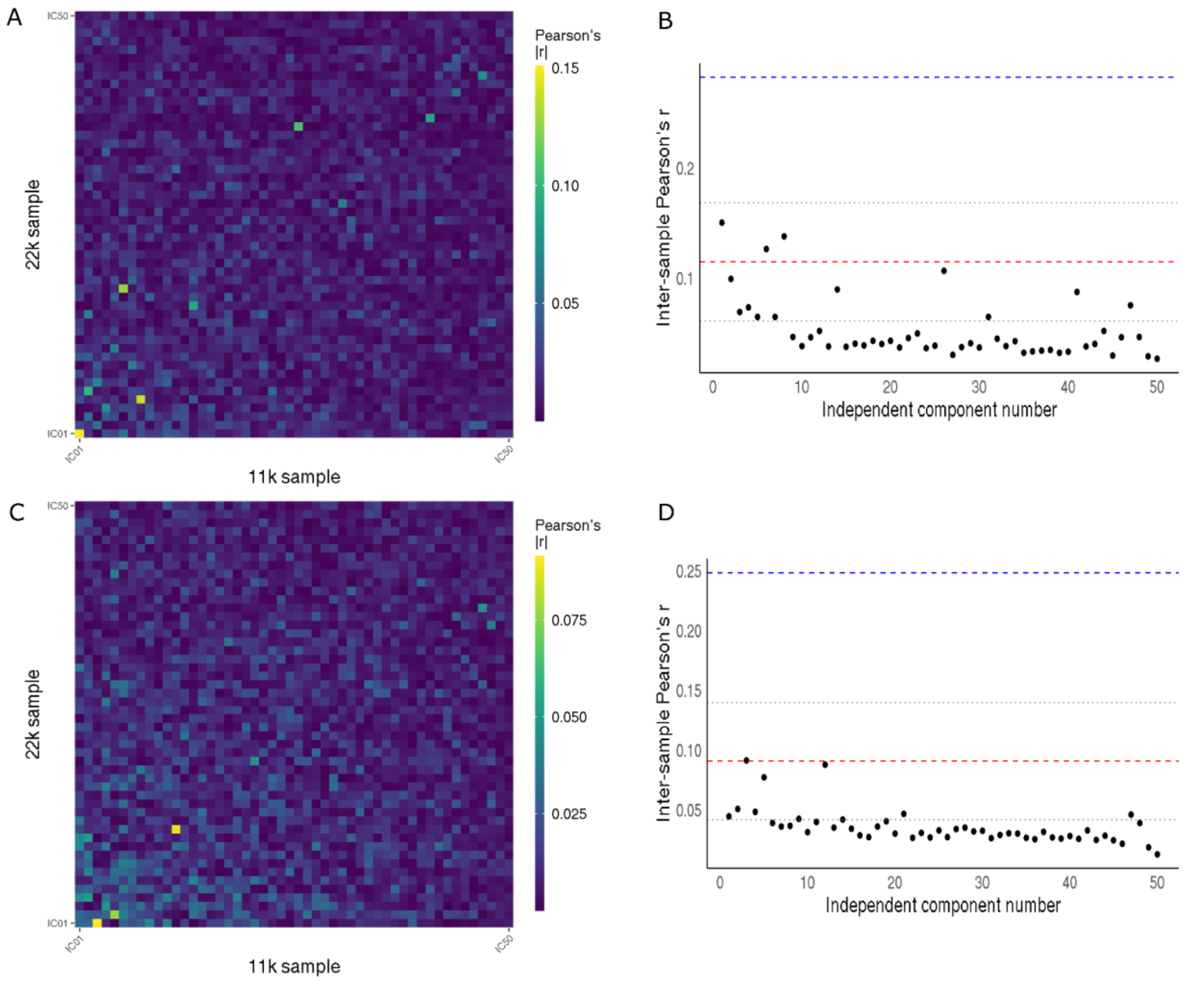


**Figure S5**: Inter-sample reproducibility of independent genomic components derived at dimension 50, from z-transformed (top row) and raw (bottom row), univariate GWAS SNP effects, displayed as the Pearson correlation coefficient in A and C. B and D show the respective reproducibility per independent component as a scatterplot, with the Pearson correlation coefficient on the y-axis. The red dashed line denotes the mean of raw, univariate GWAS reproducibility, with the gray, dotted lines indicating one standard deviation around the mean. The blue dashed line indicates the maximum reproducibility of z-transformed, univariate GWAS betas.

### Tables

**Table S1**: Statistics of Principal Component reproducibility at dimension 50. Depicted are the strongest correlating components between independent samples. The 22k sample is the discovery sample, the 11k sample is the replication sample. The decomposition of z-transformed GWAS is on the left-hand side, and the decomposition of raw GWAS is on the right-hand side. P-values are adjusted for MCC as specified in the methods section in the main manuscript. Calculations were made in R version 4.1.1, which rounds all p-values below $1^{-308}$ down to 0, which is why the correlation p-values are shown in this way. The 20 strongest component correlations are shown. The kurtotic components of the raw decomposition made a Fisher's test inappropriate, thus no p-values are provided there.

| **Z-transformed Decomposition** | | | | | **Raw Decomposition** | | | |
|---|---|---|---|---|---|---|---|---|
| **22k sample component** | **11k sample component** | **Coefficient (Pearson's \|r\|)** | ***Correlation p*-value** | **Fisher's *p*-value** | **22k sample component** | **11k sample component** | **Coefficient (Pearson's \|r\|)** | ***Correlation p*-value** |
| PC1 | PC1 | 0.33 | $<10^{-308}$ | $8.16^{-246}$ | PC1 | PC1 | 0.30 | $<10^{-308}$ |
| PC2 | PC2 | 0.25 | $<10^{-308}$ | $6.69^{-47}$ | PC2 | PC2 | 0.23 | $<10^{-308}$ |
| PC3 | PC3 | 0.26 | $<10^{-308}$ | $1.15^{-38}$ | PC3 | PC3 | 0.24 | $<10^{-308}$ |
| PC4 | PC4 | 0.21 | $<10^{-308}$ | $1.19^{-35}$ | PC4 | PC4 | 0.17 | $<10^{-308}$ |
| PC5 | PC5 | 0.18 | $<10^{-308}$ | $1.22^{-28}$ | PC5 | PC5 | 0.13 | $<10^{-308}$ |
| PC6 | PC6 | 0.16 | $<10^{-308}$ | $2.95^{-19}$ | PC6 | PC6 | 0.11 | $<10^{-308}$ |
| PC7 | PC7 | 0.17 | $<10^{-308}$ | $5.30^{-19}$ | PC7 | PC7 | 0.13 | $<10^{-308}$ |
| PC8 | PC8 | 0.15 | $<10^{-308}$ | $1.31^{-23}$ | PC8 | PC8 | 0.10 | $<10^{-308}$ |
| PC9 | PC9 | 0.13 | $<10^{-308}$ | 0.03 | PC9 | PC9 | 0.08 | $2.17^{-256}$ |
| PC10 | PC10 | 0.12 | $<10^{-308}$ | $5.14^{-08}$ | PC14 | PC14 | 0.08 | $3.22^{-220}$ |
| PC13 | PC13 | 0.11 | $<10^{-308}$ | 0.003 | PC12 | PC12 | 0.08 | $1.84^{-202}$ |
| PC14 | PC14 | 0.12 | $<10^{-308}$ | 0.0001 | PC13 | PC13 | 0.07 | $1.69^{-199}$ |
| PC15 | PC15 | 0.11 | $<10^{-308}$ | 0.0077 | PC10 | PC10 | 0.07 | $1.13^{-196}$ |
| PC20 | PC20 | 0.09 | $8.22^{-306}$ | >0.05 | PC15 | PC15 | 0.07 | $7.60^{-194}$ |
| PC19 | PC19 | 0.09 | $3.79^{-288}$ | >0.05 | PC11 | PC11 | 0.06 | $1.80^{-135}$ |
| PC18 | PC18 | 0.09 | $7.33^{-267}$ | >0.05 | PC17 | PC16 | 0.05 | $6.99^{-107}$ |
| PC16 | PC17 | 0.08 | $7.30^{-260}$ | >0.05 | PC19 | PC19 | 0.05 | $2.51^{-100}$ |
| PC17 | PC16 | 0.08 | $1.25^{-243}$ | >0.05 | PC22 | PC22 | 0.05 | $3.25^{-90}$ |

| | | | | | | | | |
|---|---|---|---|---|---|---|---|---|
| PC24 | PC23 | 0.07 | $2.80^{-177}$ | >0.05 | PC20 | PC20 | 0.05 | $1.56^{-87}$ |

**Table S2**: Statistics of Independent Component reproducibility at dimension 10. Depicted are the strongest correlating components between independent samples. The 22k sample is the discovery sample, the 11k sample is the replication sample. The decomposition of z-transformed GWAS is on the left-hand side, and the decomposition of raw GWAS is on the right-hand side. P-values are adjusted for MCC as specified in the methods section in the main manuscript. Calculations were made in R version 4.1.1, which rounds all p-values below $1^{-308}$ down to 0, which is why the correlation p-values are shown in this way. The 20 strongest component correlations are shown. The kurtotic components of the raw decomposition made a Fisher's test inappropriate, thus no p-values are provided there.

| **Z-transformed Decomposition** | | | | | **Raw Decomposition** | | | |
|---|---|---|---|---|---|---|---|---|
| **22k sample component** | **11k sample component** | **Coefficient (Pearson's \|r\|)** | ***Correlation p*-value** | **Fisher's *p*-value** | **22k sample component** | **11k sample component** | **Coefficient (Pearson's \|r\|)** | ***Correlation p*-value** |
| IC1 | IC2 | 0.23 | $<10^{-308}$ | $5.47^{-66}$ | IC1 | IC1 | 0.19 | $<10^{-308}$ |
| IC6 | IC6 | 0.20 | $<10^{-308}$ | $1.01^{-11}$ | IC2 | IC1 | 0.09 | $<10^{-308}$ |
| IC9 | IC9 | 0.15 | $<10^{-308}$ | $7.30^{-07}$ | IC7 | IC2 | 0.13 | $<10^{-308}$ |
| IC7 | IC7 | 0.14 | $<10^{-308}$ | >0.05 | IC2 | IC3 | 0.10 | $<10^{-308}$ |
| IC10 | IC10 | 0.12 | $<10^{-308}$ | 0.003 | IC4 | IC3 | 0.12 | $<10^{-308}$ |
| IC2 | IC5 | 0.11 | $<10^{-308}$ | >0.05 | IC4 | IC4 | 0.10 | $<10^{-308}$ |
| IC5 | IC7 | 0.10 | $<10^{-308}$ | >0.05 | IC3 | IC5 | 0.16 | $<10^{-308}$ |
| IC2 | IC3 | 0.11 | $<10^{-308}$ | >0.05 | IC8 | IC6 | 0.10 | $<10^{-308}$ |
| IC8 | IC3 | 0.12 | $<10^{-308}$ | >0.05 | IC6 | IC7 | 0.19 | $<10^{-308}$ |
| IC5 | IC5 | 0.13 | $<10^{-308}$ | 0.005 | IC5 | IC8 | 0.09 | $7.98^{-305}$ |
| IC3 | IC1 | 0.17 | $<10^{-308}$ | >0.05 | IC2 | IC4 | 0.09 | $1.72^{-293}$ |
| IC4 | IC4 | 0.17 | $<10^{-308}$ | >0.05 | IC4 | IC9 | 0.08 | $4.48^{-203}$ |
| IC8 | IC8 | 0.18 | $<10^{-308}$ | >0.05 | IC2 | IC2 | 0.07 | $2.44^{-183}$ |
| IC7 | IC5 | 0.09 | $3.75^{-296}$ | >0.05 | IC4 | IC10 | 0.07 | $9.11^{-174}$ |
| IC2 | IC4 | 0.09 | $1.58^{-276}$ | >0.05 | IC9 | IC9 | 0.07 | $1.34^{-171}$ |
| IC4 | IC3 | 0.09 | $2.20^{-275}$ | >0.05 | IC8 | IC4 | 0.07 | $1.48^{-164}$ |
| IC1 | IC5 | 0.08 | $2.10^{-257}$ | >0.05 | IC1 | IC3 | 0.07 | $6.76^{-160}$ |
| IC4 | IC6 | 0.08 | $5.23^{-226}$ | >0.05 | IC9 | IC7 | 0.07 | $7.95^{-158}$ |

| | | | | | | | | |
|---|---|---|---|---|---|---|---|---|
| IC2 | IC8 | 0.07 | $3.05^{-196}$ | >0.05 | IC5 | IC6 | 0.07 | $1.88^{-157}$ |
